# 3D label-free imaging and analysis of *Pinus* pollen grains using optical diffraction tomography

**DOI:** 10.1101/219378

**Authors:** Geon Kim, SangYun Lee, Seungwoo Shin, YongKeun Park

## Abstract

- The structure of pollen grains is related to the reproductive function of the plants. Here, three-dimensional (3D) refractive index maps were obtained for individual conifer pollen grains using optical diffraction tomography (ODT).
- The 3D morphological features of pollen grains from pine trees were investigated using measured refractive index maps, in which distinct substructures were clearly distinguished and analyzed.
- Morphological and physiochemical parameters of the pollen grains were quantified from the obtained refractive index (RI) maps and used to quantitatively study the interspecific differences of pollen grains from different strains.
- Our results demonstrate that ODT can assess the structure of pollen grains. This label-free and rapid 3D imaging approach may provide a new platform for understanding the physiology of pollen grains.

## Introduction

Pollen grains are the male gametophytes of seed plants and are essential to the life cycle of many plants that play a major role in ecosystems. The study of pollen also has a key role in multiple fields of science. For instance, many crucial perspectives, ranging from evidence of geological history in paleontology (Lewin, 1984), to plant reproduction (Knight *et al*., 2005), and to breeding technology in agriculture (Shivanna & Sawhney, 1997), have been advanced by investigations of pollen grains. However, measuring the three-dimensional (3D) structure of individual pollen grains remains challenging, mainly due to the limitations of conventional imaging approaches. Many conventional imaging methods lack the capacity for quantitative 3D measurement, and even advanced 3D imaging methods have limitations or may require highly specialized equipment to image pollen grains, particularly those of conifer plants. Bright field microscopy and electron microscopy have been conventionally used for imaging pollen grains, yet they have distinct limitations. Bright field microscopy allows convenient real-time observation of pollen grains, yet the resulting images have only provided qualitative 2D information (deWin *et al*., 1996; Derksen *et al*., 1999). For high-resolution imaging of conifer pollen grains, electron microscopy has been employed. In previous studies, scanning electron microscopy has provided detailed images of *Pinus* pollen grains (Bagnell, 1975; Runions *et al*., 1999; Bykowska & Klimko, 2015; Shi *et al*., 2015). However, scanning electron microscopy can only measure the surface attributes of pollen grains that have been prepared in advance using processes such as coating with gold or chromium.

In another study, transmission electron microscopy was employed to study sections of pollen grains of fixed conifer plants (Runions *et al*., 1999). However, transmission electron microscopy is not capable of measuring entire pollen grains because the specimens are restricted to slices of nanometer-scale thickness, while whole pollen grains are up to tens of micrometers.

3D optical imaging methods have also been exploited to acquire volumetric images of conifer pollen grains. For instance, fluorescence imaging can provide 3D images of fluorescent molecules in conifer pollen grains. In a previous study, 3D imaging of autofluorescence signals from conifer pollen grains was performed (Punyasena *et al*., 2012). Studies which imaged conifer pollen grains using an external dye (Anderhag *et al*., 2000) and transgenic fluorescent protein (Tian *et al*., 1997) also suggest that various molecules can be targeted to enhance the 3D imaging of conifer pollen grains. However, fluorescence imaging may chemically alter the sample (Dixit & Cyr, 2003), produce photobleaching (Song *et al*., 1997), or interfere with the intrinsic pigments of the sample (Zhou *et al*., 2005). It is also noteworthy that fluorescence images can only provide information about the distribution of fluorescent molecules.

Recently, tomographic imaging of *Pinus* pollen grains was demonstrated using a hard X-ray source (Li *et al*., 2016). However, the utilization of hard X-rays creates the risk of radiation damage to the sample, and studies may be hampered by limited access to appropriate X-ray imaging facilities.

Lately, quantitative phase imaging (QPI) methods have emerged, and because they offer quantitative and label-free imaging capability, now play an important role in the study of cellular physiologies (Popescu, 2011; Lee *et al*., 2013). By exploiting the principle of laser interferometry, QPI can quantitatively measure the phase delay of light scattered by a sample. QPI does not rely on an exogenous labeling process, because the phase delay, which is equivalent to the multiplied product of the refractive index (RI) and the thickness of the sample, is determined by the inherent composition of the sample (Kemper & von Bally, 2008).

More recently, 3D QPI techniques including optical diffraction tomography (ODT) have been extensively utilized for imaging and analyzing the internal 3D structures of various transparent biological samples, including yeast cells (Habaza *et al*., 2015), mammalian blood cells (Kim, K *et al*., 2014; Hur *et al*., 2017; Lee, S *et al*., 2017; Merola *et al*., 2017; Yoon *et al*., 2017), mammalian eukaryotic cells (Kim, T *et al*., 2014; Kim *et al*., 2016; Yang *et al*., 2017), and microorganisms (Cotte *et al*., 2013; Lee *et al*., 2014). ODT is a technique that reconstructs a 3D RI map from light scattered from a sample subjected to illumination at different angles (Wolf, 1969). The measured RI is the retardation coefficient of light speed and is related to the density of dipoles in the medium. As a result, using ODT, the structure of a sample can be identified in 3D from the reconstructed RI map.

In this study, we measured and analyzed the 3D structures of pollen grains from *Pinus* plants using ODT. Characteristic morphological features of the pollen grains were investigated using the RI maps. In addition, we quantified three parameters of individual pollen grains from their RI maps and used them to identify the morphological and chemical properties of the pollen grains.

## Materials and Methods

### Sample collection and preparation

The pollen grains were collected during late April from blooming pine trees of a local farm. Pollen grains were placed in containers by gently shaking them off the male cones of the trees. They were dried for approximately 2 hours at room temperature, then sealed and preserved in a dark cabinet at room temperature. In order to prepare pollen grains for imaging, they were suspended in a drop of RI-matching oil (Series A, Cargille Labs, United States) and sandwiched between two cover glasses (C024501, Matsunami Glass Ind. Ltd., Japan). Imaging was conducted using pollen grains that were not aggregated with other particles.

### Image acquisition and 3D RI map reconstruction

Using ODT, the 3D RI map of a sample was reconstructed using holographic images of a sample that had been illuminated from different angles (see Fig. 1a for the schematic diagram). The acquisition of phase and intensity images was performed using an off-axis Mach-Zehnder interferometric setup with a digital micromirror device (DMD) installed (HT-1S, Tomocube Inc., Republic of Korea) (see Fig. 1b). The incident angles of illumination were rapidly scanned by projecting a series of binary holograms on the installed DMD (Shin *et al*., 2015; Lee, K *et al*., 2017). The light scattered by the sample was transmitted to the camera along an imaging system consisting of an objective lens (60×, numerical aperture = 0.8) and a tube lens *f* = 175 mm). On the camera plane, the transmitted light interferes with a plane reference beam, which is obliquely incident on the camera. The spatial resolution of the microscope is 166 nm and 1 μm for lateral and axial direction, respectively. For a single pollen grain, the measurement of scattered fields required less than a second.

**Figure 1.**
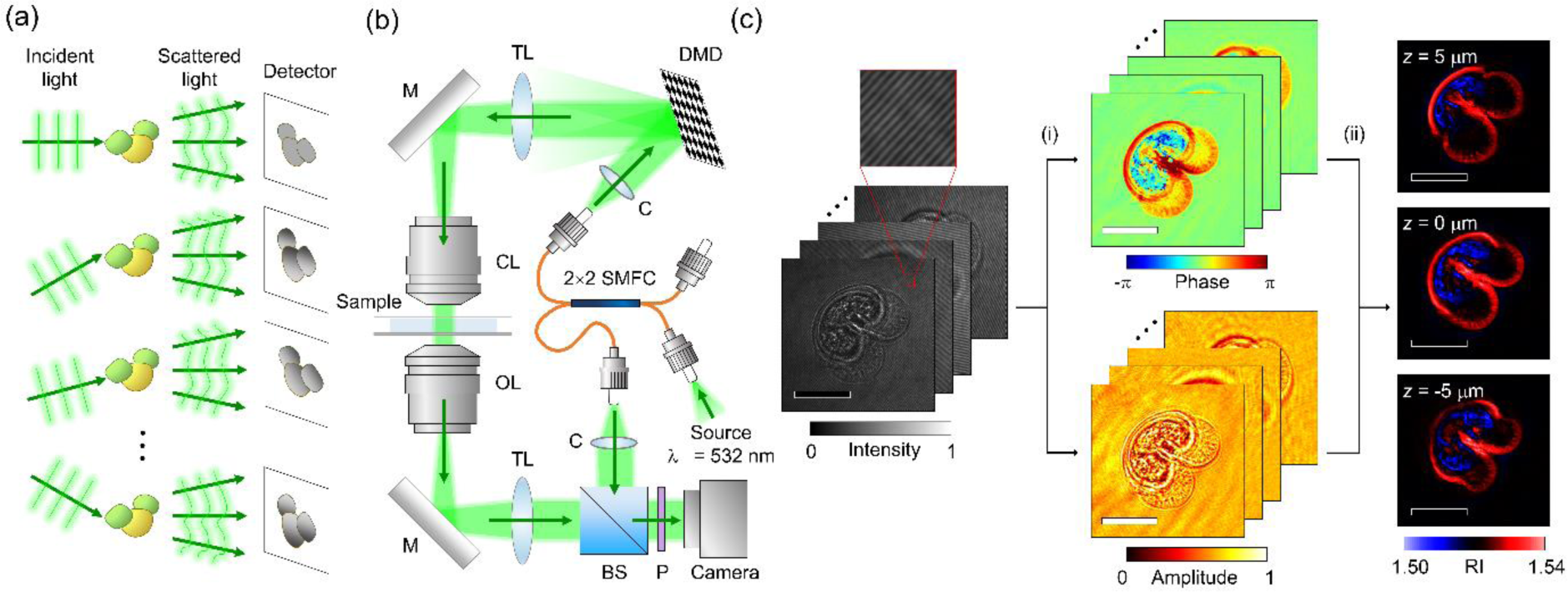
Reconstruction of a three-dimensional (3D) refractive index (RI) map using optical diffraction tomography. (a) A schematic diagram of illumination angle scanning. (b) The optical setup for measurement of the light scattered by the sample. 2×2 SMFC: 2-by-2 single-mode fiber optics coupler; C: collimator; TL: tube lens; M: mirror, CL: condenser lens; OL: objective lens, BS: beam splitter; P: polarizer, (c) Image processing for reconstruction of 3D RI maps from angle-scanned holograms. (i) indicates the optical field retrieval process, and (ii) indicates the tomographic reconstruction process. Scale bar = 20 μm.

The phase and amplitude of the scattered light can be retrieved from the recorded interference pattern, and then processed for reconstruction of the 3D RI map (see Fig. 1c). Reconstruction of the RI maps was conducted using the principle previously proposed (Wolf, 1969). Detailed descriptions of the reconstruction algorithm can be found elsewhere (Kim, K *et al*., 2014; Lim *et al*., 2015).

Using ODT, 3D RI maps of a total of 92 entire pollen grains and 175 hollow shells of pollen grains were reconstructed. From Korean red pine trees *(Pinus densiflora)*, a total of 6 pollen grains and 113 shells were imaged. From golden Korean red pine trees *(Pinus densiflora Aurea)*, 17 pollen grains and 37 shells were imaged. From Japanese red pine trees *(Pinus densiflora for. multicaulis)*, 38 pollen grains and 12 shells were imaged. From Japanese black pine trees *(Pinus thunbergii)*, 31 pollen grains and 13 shells were imaged.

## Result

### 3D structures of *Pinus* pollen grains in RI maps

RI maps of individual pollen grains from four types of local *Pinus* plants were reconstructed using ODT. Analogous to the sectional images obtained in X-ray computed tomography, 3D RI maps can be visualized at lateral cross-sections (see Fig. 2). The bisaccate morphology, which is a commonly shared trait of pollen grains from conifer plants, was clearly visible in the cross-sectional images. Due to the high level of morphological variations between individual pollen grains, an abstract description of the morphology alone is not sufficient to determine the strain of a given pollen grain.

**Figure 2.**
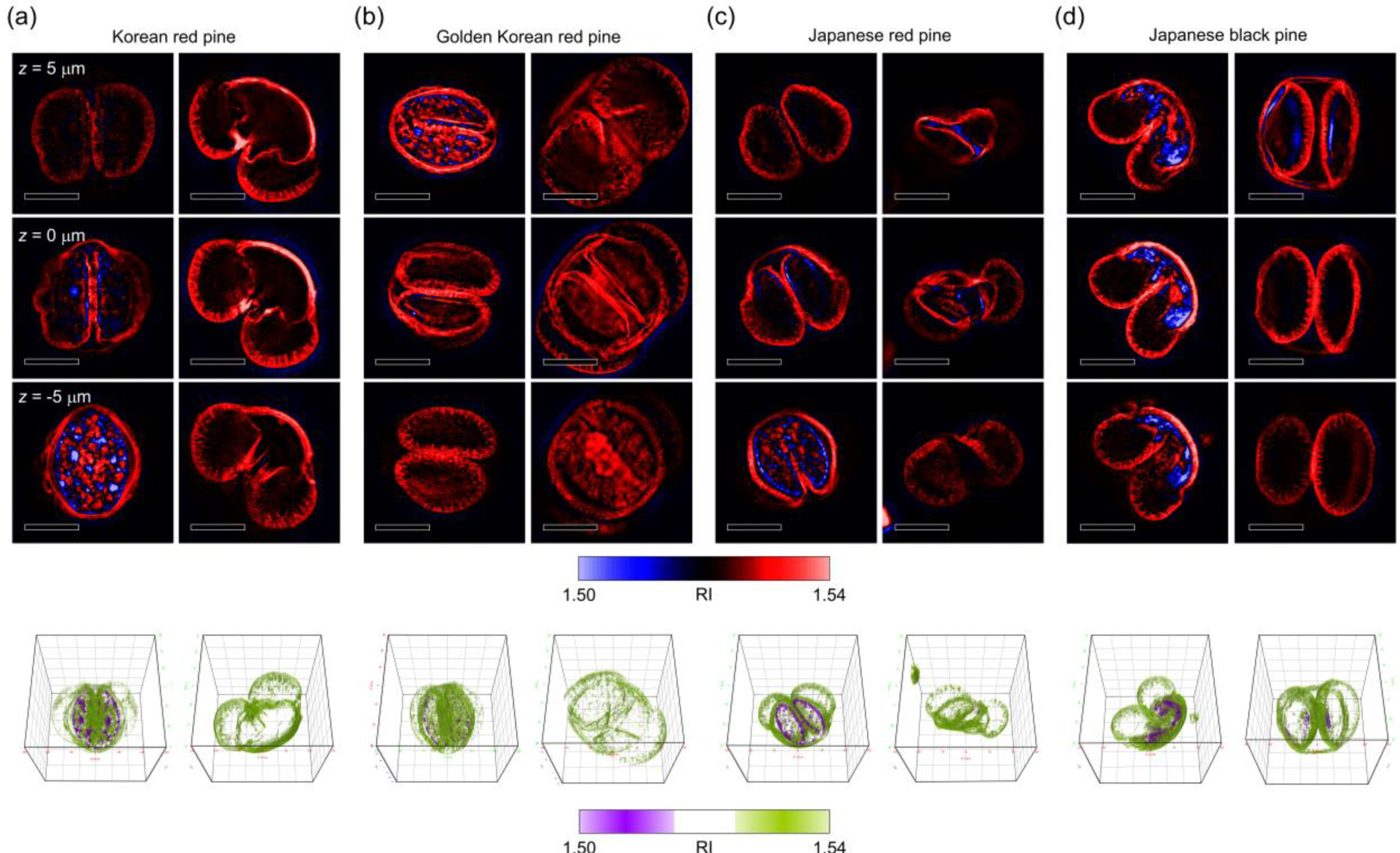
Three-dimensional (3D) refractive index (RI) maps of individual *Pinus* pollen grains. Axial section images (upper) and 3D rendering images (lower) of pollen grains from Korean red pine (a), Korean golden pine (b), Japanese red pine (c), and Japanese black pine. For each strain, the left column and the right column presents a whole pollen grain and an empty shell, respectively. Three upper three rows display the sectional RI images at different axial depths. In the lower row, RI between 1.5 and 1.515 is rendered in violet and RI between 1.525 and 1.54 is rendered in yellow-green. Scale bar = 20 μm. Cube length = 60 μm.

In order to demonstrate the 3D imaging capability of ODT, detailed substructures of *Pinus* pollen grains were further investigated. Structures of exines, or integumentary shells of pollen grains, could be observed in the RI maps of the hollow shells of pollen grains (see Fig. 3a). The shell of a pollen grain was distinguished from the medium in the RI map. The shell exhibited RI over 1.53 which was 0.01 higher than the medium RI. The distinction between the medium and the shell in RI provided a sharp outline of each pollen grain.

**Figure 3.**
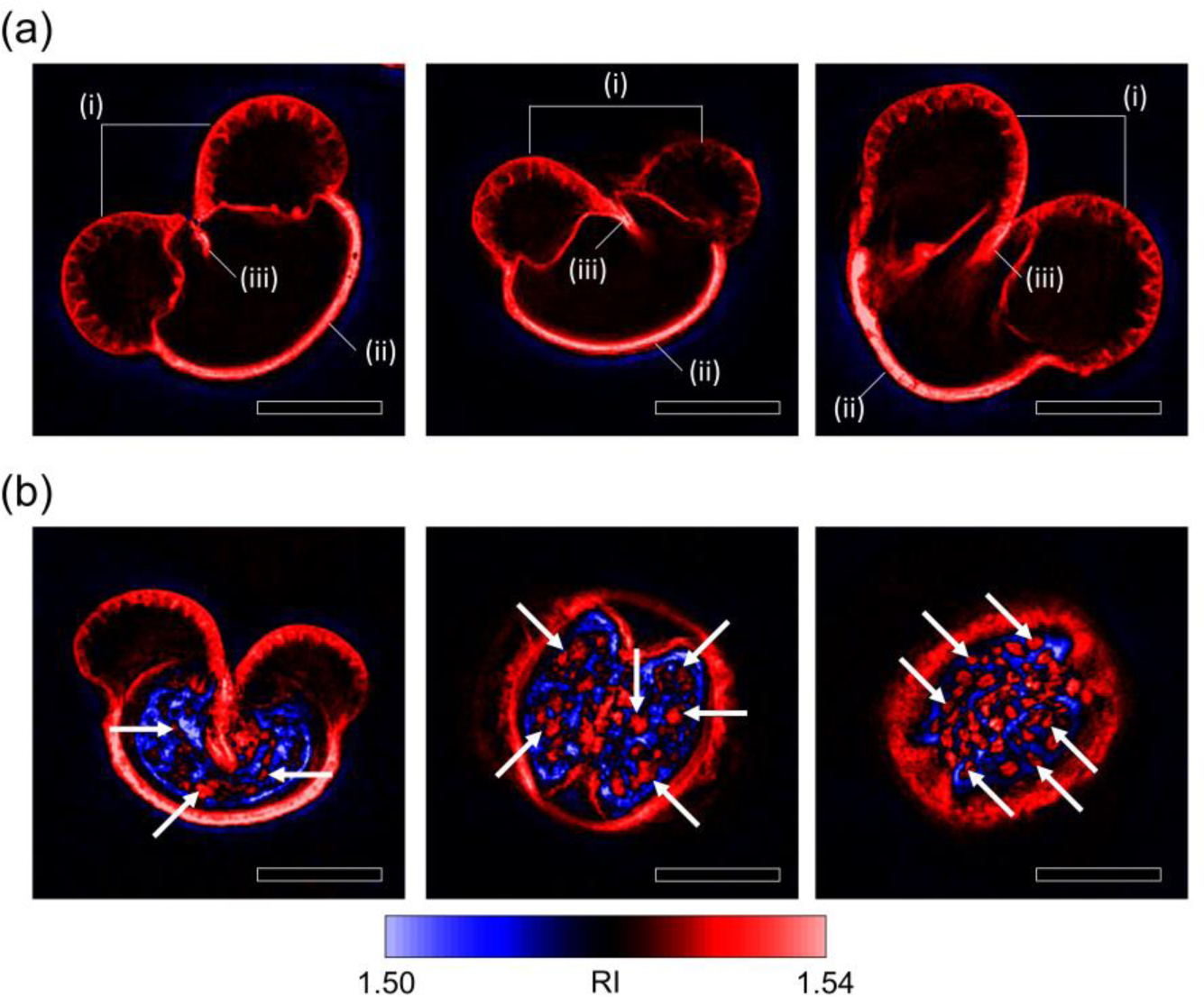
Characteristic structures of pine pollen grains. (a) Axial section images of the hollow shells of pollen grains that exhibit distinguishing exine structures. The sacci (i) are shown to be empty and macroporous exine sacks of refractive index (RI) over 1.53. The cappa (ii) appears as a thick and rigid exine wall of RI near 1.54. The germinal wrinkles (iii) are located at the ventral side of a pollen grain between the two sacci. (b) Axial section images of whole pollen grains. Starch granules (given examples using white arrows) are visualized due to the RI contrast between starch and the rest of the corpus. Scale bar = 20 μm.

The sacci, which are air sacks attached to the pollen grains, were recognizable as paired chambers bulging towards the outside of the pollen grain (Fig. 3a; see the mark (i)). A pair of sacci was separated from the body of the pollen grain with a thin exine screen, while the outer surfaces were composed of the thick yet macroporous walls of the exine. The sacci were depicted as filled with the background medium since the RI level inside the sacks was highly homogeneous near the medium RI.

The cappa, which refers to the thick region of an exine at the dorsal pole of a pollen grain (Bagnell, 1975), was visualized in the RI maps as a robust and smooth exine barrier with an RI near 1.54 (Fig. 3a; see the mark (ii)). The RI map showed that the cappa of a *Pinus* pollen grain covers a wide area of the dorsal exine. Wrinkles at the ventral poles of the exine shells were depicted in the RI maps as ravines in the exine shells between the air sacks (Fig. 3a; see the mark (iii)). The ends of the wrinkled exine were penetrating towards the centers of the pollen grains; this is an observation which was not revealed in scanning electron microscopy images of conifer pollen grains (Bagnell, 1975; Runions *et al*., 1999).

The internal substructures of the pollen grains were investigated using RI maps of the entire pollen grains (see Fig. 3b). Unlike the image of a hollow exine shell, the region inside the body of a whole pollen grain, which is occasionally referred to as the corpus, displayed an inhomogeneous distribution of RI. In the region, RI levels both higher and lower than the medium RI were observed. Inside the exine of a whole pollen grain, were multiple granules of 2-5 μm diameter with RI near 1.53. (Fig. 3b; see the white arrows) The granules were found to be embedded in a background of RI lower than the medium RI. The elliptical morphology, the size, and the RI levels of the granules suggested that they were starch granules stored in the pollen grains (Wolf *et al*., 1962; Fernando *et al*., 2005).

### Quantification of parameters using RI maps

Morphological parameters including volume and surface area were quantified by distinguishing the pollen grains from the medium in the RI maps (See Fig. 4a and Fig 4b). The volumes of pollen grains were 33.9 ± 10.6, 43.0 ± 14.4, 21.9 ± 5.5, and 37.4 ± 8.2 pl (mean ± SD) for Korean red pine, golden Korean red pine, Japanese red pine and Japanese black pine, respectively. The results of the Mann-Whitney test showed that there were all significant differences (P-value < 0.05) in the volumes of pollen grains between different strains of *Pinus* plants, except for between golden Korean red pine and Japanese black pine. Surface areas of the pollen grains from the plants in the same order were 7490 ± 1590, 8600 ± 2030, 5230 ± 760, and 7700 ± 1280 μm^2^. According to the Mann-Whitney test, the interspecific differences of the surface areas of pollen grains were significant except for between Korean red pine and Japanese black pine. The sphericity index was also quantified for each pollen grain (See Fig. 4c). The sphericity index is an indicator for the resemblance of a shape to a sphere. For an individual pollen grain, the sphericity index was calculated from the volume and the surface area of the pollen grain, using the following formula: *SI=(36nV^2^)^13^/S*, where *SI, V* and *S* indicate the sphericity index, the volume, and the surface area of the pollen grain respectively. The sphericity indices of the pollen grains from the plants of the identical order were 0.674 ± 0.078, 0.686 ± 0.077, 0.720 ± 0.053, and 0.703 ± 0.058. In the result of the Mann-Whitney test, sphericity indices of pollen grains were significantly different between Korean red pine and Japanese red pine, and between **Korean red pine and Japanese black pine**. In addition, information about the distribution of chemical components in pollen grains was quantified from the RI maps of entire pollen grains. Masses of starch, which is a major source of nutrition in pollen grains, were quantified for individual pollen grains (See Fig. 4d). Starch, which corresponds to high-RI granules inside the exines of pollen grains, was specifically identified and investigated. Using the density of dry starch, the masses of starch could be quantified. The starch contents were 1.007 ± 1.483, 1.193 ± 1.327, 0.894 ± 0.535, and 0.171 ± 1.671 ng, respectively for pollen grains from Korean red pine, golden Korean red pine, Japanese red pine, and Japanese black pine. Due to the high variation between individual pollen grains, the standard deviation of starch content was comparable to the mean value in every strain we studied. The results of the Mann-Whitney test indicated there was no significant difference in starch content among the experimentally tested strains of *Pinus* plants.

**Figure 4.**
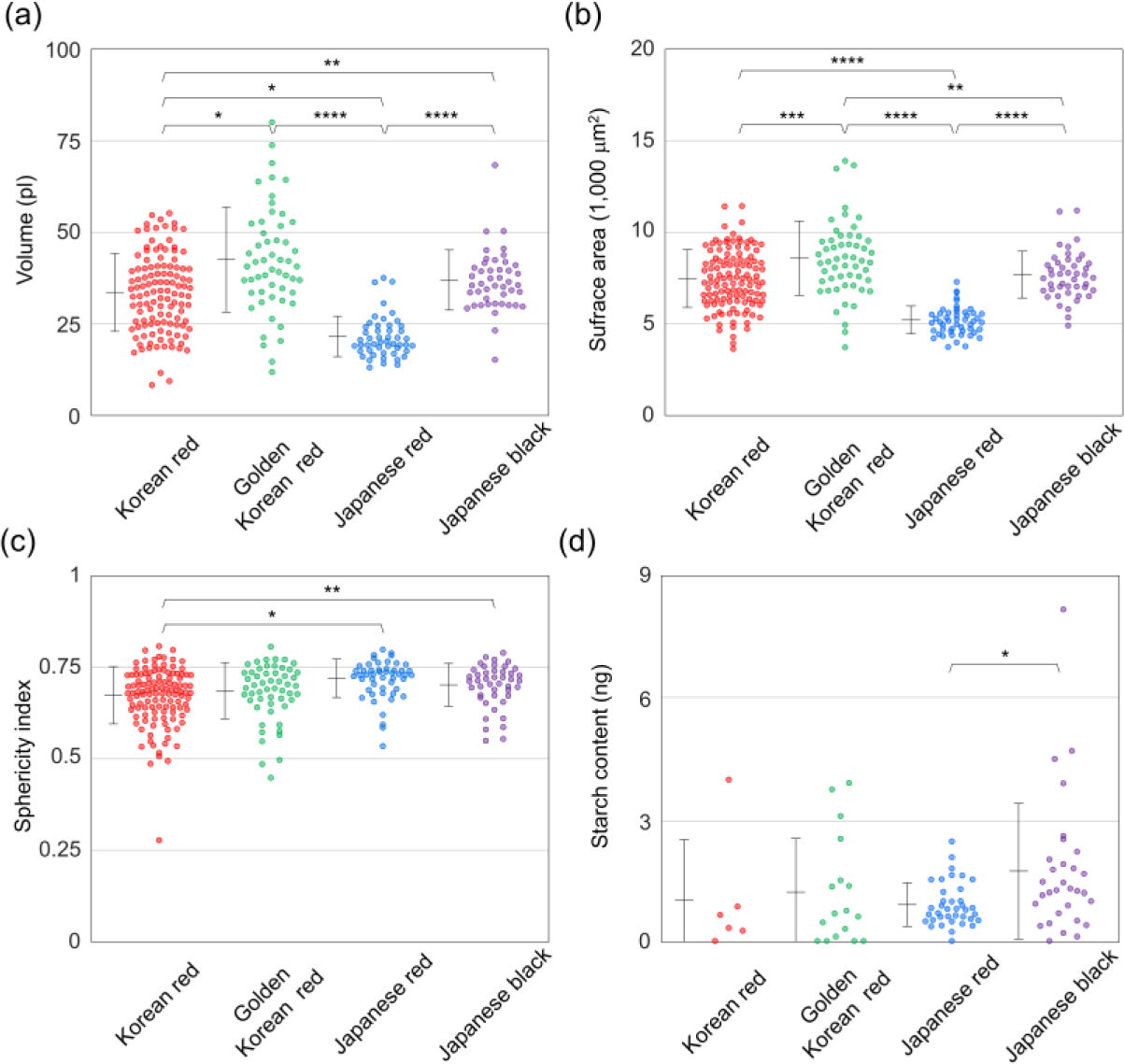
Quantification of parameters from pollen grains. (a) The volumes, (b) surface areas, (c) sphericity indices, and (d) starch contents of individual pollen grains, quantified from the 3D RI maps. Each error bars placed on the left of the scatter plots represent mean ± SD. *, **, and *** refers to *P*-value less than 0.05, 0.01, and 0.001, respectively in the Mann-Whitney U-test. The absence of an asterisk refers to difference of no significance in the Mann-Whitney U-test.

## Discussion

To summarize, we demonstrated and obtained label-free imaging and analysis of the 3D structures of *Pinus* pollen grains. From the reconstructed 3D RI maps obtained using ODT, detailed morphological features of individual *Pinus* pollen grains were identified. Furthermore, quantitative parameters which represent the structures of the pollen grains were quantified using the maps to highlight the variances within and between pollen grains from different strains of *Pinus* plants.

We expect that the study of pollen grains using ODT will illuminate previously unreachable aspects of pollen grains, with the aid of further supplementation. While 3D imaging is essential to understanding the microscopic structures of pollen grains, previous imaging techniques have either lacked 3D imaging capabilities or introduced issues of sample deterioration and poor accessibility. In our demonstration, ODT delivered high-resolution 3D images of *Pinus* pollen grains label-free. ODT also outperformed previous imaging techniques with respect to non-invasiveness and acquisition rate, with a relatively simple setup. These ODT advantages are important features in a tool, especially for studying dynamic processes in pollen grains, including germination.

While the aforementioned strengths of ODT are remarkable, the technique does not presently allow identification with high chemical specificity, although this limitation can be further improved using other techniques. While RI is an inherent optical property of matter, it cannot be used to determine the chemical composition of a sample alone. Clarifying the relationships between the structures in RI maps and their chemical constituents can be accomplished by inter-calibrating the ODT technique with other techniques that provide high chemical accuracy, including chemically resolving assays and image registration with fluorescence imaging (Schmidt & Herman, 2008; Punyasena *et al*., 2012). Advances in high-modality QPI techniques may also provide additional chemical information without labeling. Recently, a method for hyperspectral ODT was developed, demonstrating the reconstruction of multiple RI maps with various wavelengths for illumination (Jung *et al*., 2016). Since plant tissues have distinct absorption spectra, measuring complex RI depending on frequency can provide additional information that can be used to localize the absorptive components in pollen grains.

Birefringence is another property that can be exploited to characterize pollen grains. Crystalline starch, which is one of the nutritional sources in pollen grains, can be detected using birefringence (Franchi *et al*., 1996), and the measurement of full Jones matrices of microscopic samples has been demonstrated, using quantitative phase imaging and polarization filtering (Park *et al*., 2014).

We expect ODT to provide a new platform for understanding the physiology of pollen grains. For example, the time-lapse 3D imaging of pollination or pollen tube growth may enable to investigate new physiological studies (Edlund *et al*., 2004; Chae & Lord, 2011). This rapid and label-free method of 3D imaging can be further enhanced to resolve chemical components using various methods. Since rich information about the reproduction of plants lies in the structures of pollen grains, ODT will be a promising tool for understanding the life cycles of plants.

## Acknowledgements

This work was supported by KAIST, BK21+ program, Tomocube Inc., and National Research Foundation of Korea (2015R1A3A2066550, 2017M3C1A3013923, 2014K1A3A1A09063027). Mr. Shin and Prof. Park have financial interests in Tomocube Inc., a company that commercializes ODT and QPI instruments and is one of the sponsors of the work. Author Contribution: G.K. and Y.K.P. designed the study. G.K. performed the experiments and analyzed data. S.Y.L. and S.S. provide analysis methods and analyzed data. All authors wrote the manuscript.

